# *De novo* biosynthesis of *para*-nitro-L-phenylalanine in *Escherichia coli*

**DOI:** 10.1101/2021.09.29.462267

**Authors:** Neil D. Butler, Sabyasachi Sen, Minwei Lin, Aditya M. Kunjapur

## Abstract

Nitroaromatic functional groups can impart valuable properties to chemicals and to biological macromolecules including polypeptides. *Para*-nitro-L-phenylalanine (pN-Phe) is a nitroaromatic amino acid with uses including immune stimulation and fluorescence quenching. As the chemical synthesis of pN-Phe does not follow green chemistry principles and impedes provision of pN-Phe to engineered bacterial cells in some contexts, we sought to design a *de novo* biosynthetic pathway for pN-Phe in *Escherichia coli*. To generate the nitro chemical functional group, we identified natural diiron monooxygenases with measurable *in vitro* and *in vivo* activity on envisioned amine-containing precursors of *para*-amino-L-phenylalanine (pA-Phe) and *para*-aminophenylpyruvate. By expressing one of these *N*-oxygenase genes together with previously characterized genes for the biosynthesis of pA-Phe, we achieved the synthesis of pN-Phe from glucose. Through further optimization of the chassis, plasmid constructs, and media conditions, we were able to improve the selectivity of pN-Phe biosynthesis, resulting in a maximum titer of 819 µM in rich defined media under shake-flask conditions. These results provide a foundation for the biosynthesis of related nitroaromatic chemicals and for downstream biological applications that could utilize pN-Phe as a building block.

**Highlights:**

- *Para*-nitro-L-phenylalanine (pN-Phe) is a valuable small molecule for its applications in genetic code expansion.
- We establish *de novo* biosynthesis of pN-Phe from glucose in *E. coli*, which is also the first example of a *de novo* pathway design for an unnatural but commonly used non-standard amino acid.
- We show the first use of an *N*-oxygenase enzyme in the *de novo* synthesis of a nitroaromatic product.
- Screening of natural *N-*oxygenases and strain engineering resulted in final pN-Phe titers of 820 ± 130 µM in shake flask experiments with rich defined media.

## 1. Introduction

Nitro-group containing compounds, and particularly nitroaromatics, are an important class of chemicals with industrial applications as herbicides, energetic materials, and pharmaceuticals (Booth, 2000). In biological environments, the polarity of nitro-groups generates unique binding interactions with protein receptors and other molecules which enables application of nitroaromatics as antibiotics, antiparasitics, and chemotherapeutics (Nepali et al., 2019; Strauss, 1979). Specifically, the strong π electron delocalizing properties of the nitro group within aromatic systems can enhance intermolecular interactions including arene-arene and aryl hydrogen anion bonding interactions (Bryantsev and Hay, 2005; Lewis et al., 2012; Shorter, 2009). While proteins could also benefit from the installation of these properties at specific sites, none of the twenty standard amino acids exhibit similar levels of electron-withdrawing potential. For this and other reasons, researchers have been interested in using genetic code expansion technology to incorporate nitroaromatic non-standard amino acids (nsAAs) within proteins (Arbely et al., 2012; Chen et al., 2009; Deiters et al., 2006; Gautier et al., 2010; Hancock et al., 2010; Hemphill et al., 2013; Lemke et al., 2007; Neumann et al., 2008; Nguyen et al., 2014; Peters et al., 2009; Stokes et al., 2009; Tsao et al., 2006; Wang et al., 2013, 2010; Welegedara et al., 2018; Wilkins et al., 2010; Wu et al., 2004; Yanagisawa et al., 2012). The nitroaromatic non-standard amino acid *para-*nitro-L-phenylalanine (pN-Phe) in particular has been utilized in several diverse applications including as a peptide distance marker (Tsao et al., 2006), a breaker of immune self-tolerance (Gauba et al., 2011; Grünewald et al., 2009, 2008), a nitroreductase activity enhancer (Jackson et al., 2006), and an internal protein IR probe (Smith et al., 2011).

To date, the reliance on the supplementation of chemically synthesized pN-Phe has constrained the performance of cells that are engineered to harness this building block and limited their potential end-applications. Microbial biosynthesis of nsAAs presents a promising alternative because it eliminates external supplementation requirements, circumvents cellular uptake limitations, and lowers toxicity through on-demand synthesis and incorporation (Dickey et al., 2021; Völler and Budisa, 2017). Moreover, the ability to biosynthesize nsAAs from simple carbon sources should allow microbes to perform augmented protein functions autonomously in environmental and therapeutic contexts where provision is not possible (Parker and Kunjapur, 2020). Pioneering work has achieved biosynthesis of a limited number of nsAAs, including *para*-amino-L-phenylalanine (pA-Phe) (Chen et al., 2018; Mehl et al., 2003), L-dihydroxyphenylalanine (Kim et al., 2018), 4-nitro-L-tryptophan (Zuo and Ding, 2019), 5-hydroxytryptophan (Chen et al., 2020), and propargylglycine (Marchand et al., 2019). While significant advancements, these nsAAs are found in nature and were produced in model microbes by transplanting naturally occurring and highly specific pathways. To our knowledge, pN-Phe has not been found in nature (Parry et al., 2011; Waldman et al., 2017) and would thus require *de novo* pathway design for its biosynthesis.

Design of metabolic syntheses for nitroaromatic products such as pN-Phe can also provide an alternative to the harsh acidic reaction conditions and poor regioselectivity of traditional chemical nitration (Badgujar et al., 2016; Liu et al., 2009; Yan and Yang, 2013). To construct a metabolic pathway that achieves biosynthesis of pN-Phe, we anticipated that nitro group formation could proceed from aromatic metabolites via one of the two major characterized methods of enzymatic nitro synthesis: biocatalytic direct nitration or amine oxidation (Sulzbach and Kunjapur, 2020; Winkler and Hertweck, 2007). Biocatalytic direct nitration is a process dependent on two enzymatic steps: (1) the cytochrome P450-mediated nitration of aromatic rings using O2 and nitric oxide (NO) and (2) the synthesis of reactive NO from the activity of nitric oxide synthases (NOS) (Caranto, 2019). To date, only two P450/NOS pairs for aromatic direct nitration have been identified, and neither has demonstrated activity to nitrate phenylalanine (Barry et al., 2012; Dodani et al., 2016; Tomita et al., 2017; Zuo et al., 2017). Amine oxidation, on the other hand, is a promising route for pN-Phe synthesis for two reasons. First, the metabolic synthesis of the amine precursor, pA-Phe has been demonstrated from multiple natural gene clusters (commonly referred to as *papABC*) (Chen et al., 2018; Masuo et al., 2016; Mehl et al., 2003). Second, a promising amine oxidizing enzyme, or *N*-oxygenase (ObaC, otherwise referred to in the literature as ObiL), was recently discovered in *Pseudomonas fluorescens* in the biosynthetic gene cluster for the antibiotic obafluorin. Preliminary and indirect evidence suggests that this *N*-oxygenase has native activity on *para*-aminophenylpyruvate (pA-Pyr), the immediate precursor to pA-Phe in metabolic synthesis (Schaffer et al., 2017; Scott et al., 2017). Despite exploration of several *N*-oxygenases like ObaC with diiron monooxygenase activity, namely AurF (Chanco et al., 2014) (with a native substrate of para-aminobenzoic acid) and CmlI (Tsutsumi et al., 2018) (with a native substrate of the amine precursor to chloramphenicol) as potential *in vitro* catalysts for nitroaromatic product synthesis, thus far, no *N*-oxygenases have been used *in vivo* for *de novo* biosynthetic cascades despite their promise and demonstrated activity in *E. coli*.

Here, we constructed and tested a *de novo* biosynthetic pathway in *Escherichia coli* based on the following enzymes (**Fig. 1**): (i) Three enzymes from the chloramphenicol biosynthesis pathway in *Streptomyces venezuelae* that generate pA-Phe from chorismate; (ii) the putative *N*-oxygenase ObaC; (iii) the ability of native *E. coli* aminotransferases to convert keto acids to amino acids (Hayashi et al., 1993; Onuffer et al., 1995). Upon confirmation of *de novo* pN-Phe biosynthesis, we transferred the proposed enzymes to a chorismate overproducing strain of *E. coli* which further improved product titers. We then performed sequence similarity analysis to identify *N*-oxygenase enzymes related to those previously characterized and identified a candidate (NO16) that achieves higher conversion of pA-Phe than ObaC. Integration of this *N*-oxygenase improved pN-Phe titers with maximum titer observed in MOPS-EZ rich defined media.

**Figure 1.**
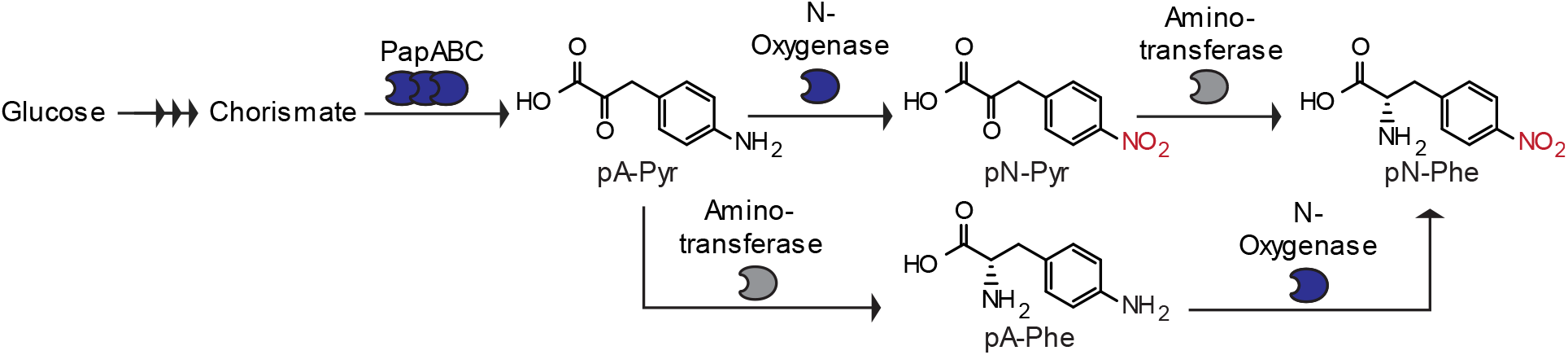
A graphical overview of the proposed *de novo* biosynthesis pathway for *para*-nitro-L-phenylalanine (pN-Phe) with exogenous enzymes shown in blue and endogenous enzymes shown in gray. PapA, 4-amino-4-deoxychorismate synthase; PapB, 4-amino-4-deoxychorismate mutase; PapC, 4-amino-4-deoxyprephenate dehydrogenase; pA-Pyr, para-aminophenylpyruvate; pN-Pyr, *para*-nitrophenylpyruvate; pA-Phe, *para*-amino-L-phenylalanine.

## 2. Materials and Methods

### 2.1. Strains and plasmids

Molecular cloning and vector propagation were performed in DH5α. Polymerase chain reaction (PCR) based DNA replication was performed using KOD XTREME Hot Start Polymerase for plasmid backbones or using KOD Hot Start Polymerase otherwise. Cloning was performed using Gibson Assembly. Genes were purchased as G-Blocks or gene fragments from Integrated DNA Technologies (IDT) or Twist Bioscience and were optimized for *E. coli* K12 using the IDT Codon Optimization Tool. The plasmids pOSIP-TH (Addgene plasmid # 45978; http://n2t.net/addgene:4598; RRID:Addgene_45988) and pE-FLP (Addgene plasmid # 45978; http://n2t.net/addgene:45978 ; RRID:Addgene_45978) were gifts from Drew Endy & Keith Shearwin (Cui and Shearwin, 2017). The *papABC* operon was kindly provided by Professor Ryan Mehl of Oregon State University in plasmid pLASC-lppPW. The pORTMAGE-EC1 recombineering plasmid was kindly provided by Timothy Wannier of the University of Ottawa (Wannier et al., 2020).

The chorismate overproducer strain was derived from a commercially available phenylalanine overproducer *E. coli* strain (NST37, ATCC #31882). To enable the compatibility of this strain with T7-promoter systems, the 4521 bp region of the phage T7 genome that is responsible for T7 polymerase functionality (*lacI, lacZ*, and T7 RNA polymerase) was genomically integrated using one-step clonetegration with the pOSIP-TH plasmid. Following plasmid assembly, NST37 was transformed with the clonetegration plasmid, and the integration of the region for T7 polymerase activity (DE3) was confirmed via selection on LB-agar plates containing 9 µg/mL tetracycline. Following Sanger sequencing-based confirmation of genetic incorporation, genomic tetracycline resistance was removed using pE-FLP. Then, using multiplexed automated genome engineering (MAGE) with the pORTMAGE-EC1 recombineering plasmid, two stop codons were introduced into the genomic sequence for the chorismate mutase/prephenate dehydratase PheA at positions 10 and 12 to serve as a translational knockout. Curing of the pORTMAGE-EC1 plasmid following Sanger sequencing confirmation of genomic knockouts produced the chorismate overproducer strain (NST37 (DE3) *ΔpheA*).

### 2.2. Chemicals

The following compounds were purchased from MilliporeSigma: hydrogen peroxide, kanamycin sulfate, chloramphenicol, carbenicillin disodium, dimethyl sulfoxide (DMSO), potassium phosphate dibasic, potassium phosphate monobasic, magnesium sulfate, calcium chloride dihydrate, imidazole, glycerol, M9 salts, sodium dodecyl sulfate, lithium hydroxide, boric acid, Tris base, glycine, HEPES, and KOD XTREME Hot Start and KOD Hot Start polymerases. pN-Phe and D-glucose were purchased from TCI America. pA-Phe, methanol, agarose, Laemmli SDS sample reducing buffer, and ethanol were purchased from Alfa Aesar. pA-Pyr and pN-Pyr were purchased from abcr GmbH. Anhydrotetracycline (aTc) and isopropyl β-D-1-thioglactopyranoside (IPTG) were purchased from Cayman Chemical. Acetonitrile, sodium chloride, LB Broth powder (Lennox), LB Agar powder (Lennox), were purchased from Fisher Chemical. Trace Elements A was purchased from Corning. L-Arabinose was purchased from VWR. A MOPS EZ rich defined medium kit was purchased from Teknova. Taq DNA ligase was purchased from GoldBio. Phusion DNA polymerase and T5 exonuclease were purchased from New England BioLabs (NEB). Sybr Safe DNA gel stain and BenchMark^™^ His-tagged Protein Standard were purchased from Invitrogen. HRP-conjugated 6*His His-Tag Mouse McAB was obtained from Proteintech. Oligonucleotides were purchases from IDT. Gene fragments were purchased from Twist Bioscience with the exception of ObaC-ObaD which was purchased from IDT.

### 2.3. Culture conditions

Cultures for general culturing, for experiments in Figures 2A, 2B, 3, 5B S4, and S6, and *N-*oxygenase protein overexpression were grown in LB-Lennox medium (LBL: 10 g/L bacto tryptone, 5 g/L sodium chloride, 5 g/L yeast extract). Cultures to demonstrate *de novo* pN-Phe synthesis were grown in either LB-Lennox-glucose medium (LBL with 1.5% glucose (wt/vol)), M9-glucose minimal media (Kunjapur et al., 2016) with Corning® Trace Elements A (1.60 µg/mL CuSO_4_ • 5H_2_O, 863.00 µg/mL ZnSO_4_ • 7H_2_O, 17.30 µg/mL Selenite • 2Na, 1155.10 µg/mL ferric citrate) and 1.5% glucose (wt/vol), or MOPS EZ rich defined media (Teknova M2105) with 1.5% glucose (wt/vol). For cultures of NST37(DE3) *ΔpheA* strains that were grown in M9-glucose minimal media, 0.04 mg/ml L-phenylalanine, 0.04 mg/ml L-tyrosine, and 0.04 mg/mL L-tryptophan were added to the media to ensure growth.

**Figure 2.**
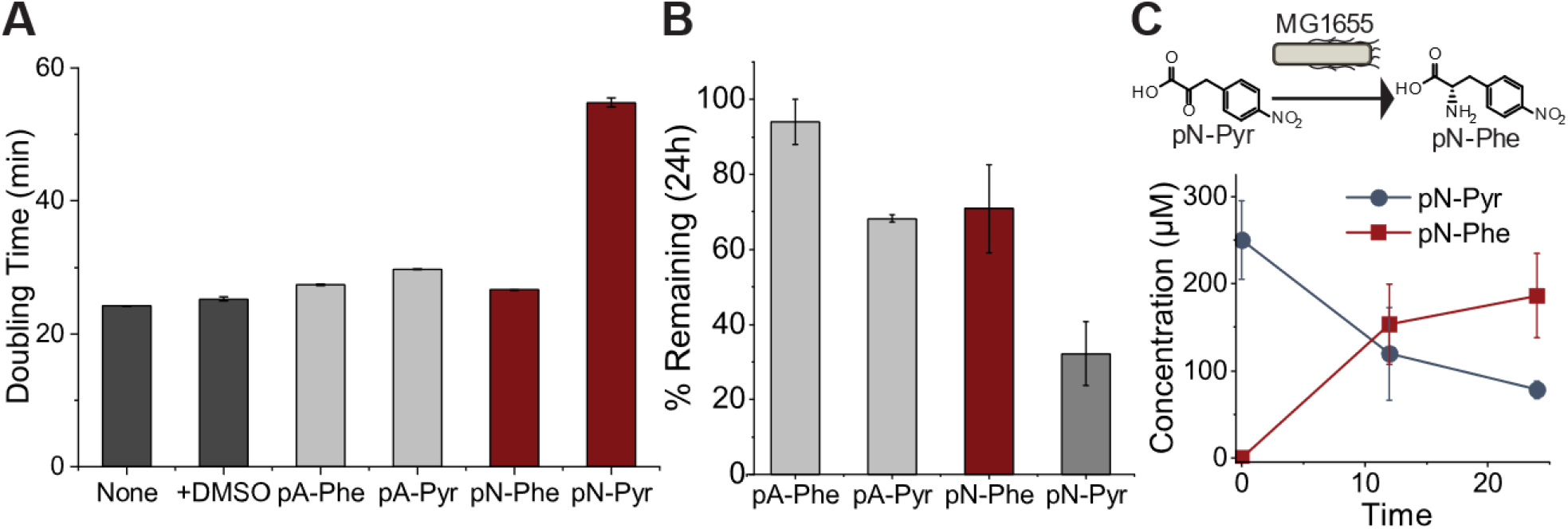
Assessment of pathway metabolite stability and toxicity. (A) Toxicity testing of pathway metabolites pA-Phe, pA-Pyr, pN-Phe, and pN-Pyr in *E. coli* MG1655(DE3) in with no additives (None), 5% DMSO for solubility (+DMSO), or 1 mM addition of associated metabolite with 5% DMSO measured via *E. coli* doubling time. (B) Stability testing of pathway metabolites measured via the percent of the initial metabolite supplementation (0.5 mM) remaining after 24 h of fermentative growth in at 37°C. (C) Tracking of pN-Pyr conversion to pN-Phe in cultures supplemented with 0.25 mM pN-Pyr over a 24 h fermentation in *E. coli* MG1655(DE3). All cultures were grown in LBL media. Sample size is *n*=3 using biological replicates. Data shown are mean ± standard deviation.

**Figure 3.**
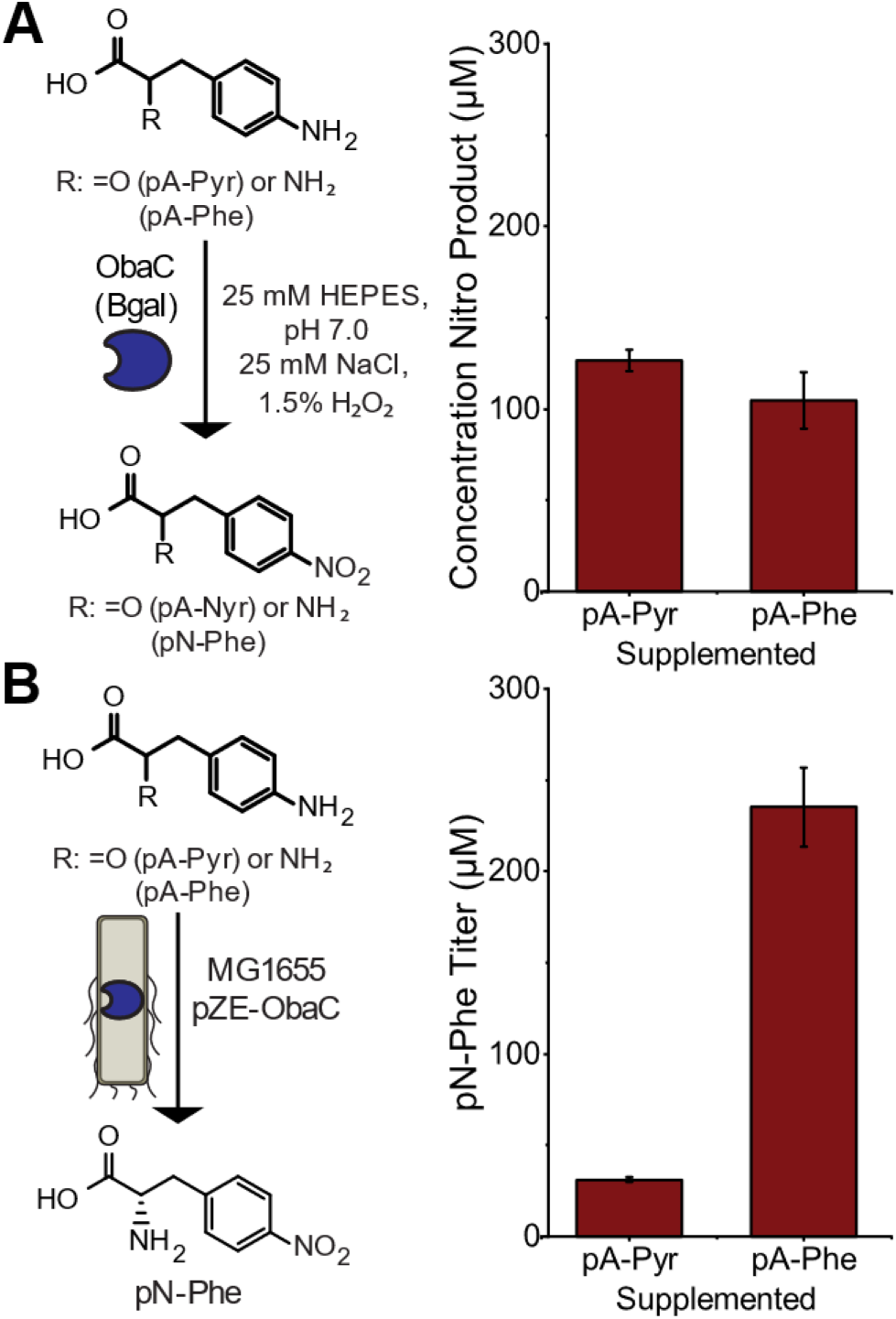
Evaluation of the activity of the *N-*oxygenase ObaC on the aromatic amines pA-Pyr and pA-Phe. (A) In vitro investigation of ObaC activity on pA-Phe and pA-Pyr. (B) *In vivo* supplementation testing of MG1655(DE3) expressing βgal-ObaC in the conversion of 2 mM supplemented pA-Pyr or pA-Phe to pN-Phe end product after 24 h of fermentative growth. Sample size is *n*=3 using biological replicates. Data shown are mean ± standard deviation.

For stability testing, a culture of *E. coli* K12 MG1655 (DE3) was inoculated from a frozen stock and grown to confluence overnight in 5 mL of LBL media. Confluent overnight cultures were then used to inoculate experimental cultures in 300 µL volumes in a 96-deep-well plate (Thermo Scientific^™^ 260251) at 100x dilution. Cultures were supplemented with 0.5 mM of heterologous metabolites (pA-Phe, pA-Pyr, pN-Pyr, pN-Phe), with pN-Pyr requiring an addition of 15 µL of DMSO (∼5% final concentration) for solubility. Cultures were incubated at 37 °C with shaking at 1000 RPM and an orbital radius of 3 mm. Compounds were quantified from the extracellular broth over a 24 h period using HPLC.

For toxicity testing, cultures were similarly prepared with confluent overnight cultures of MG1655 (DE3) used to inoculate experimental cultures at 100x dilution in 200 µL volumes in a Greiner clear bottom 96 well plate (Greiner 655090) in LBL media. Cultures were supplemented with 1 mM of heterologous metabolite and 5% DMSO for metabolite solubility and grown for 24 h in a Spectramax i3x plate reader with medium plate shaking at 37 °C with absorbance readings at 600 nm taken every 5 min to calculate doubling time and growth rate.

For supplementation testing, strains transformed with plasmids expressing pathway genes were prepared with inoculation of 300 µL volumes in a 96-deep-well plate with appropriate antibiotic added to maintain plasmids (34 μg/mL chloramphenicol (Cm), 50 μg/mL kanamycin (Kan), 50 μg/mL carbenicillin (Carb), or 95 μg/mL streptomycin (Str)). Cultures were incubated at 37 °C with shaking at 1000 RPM and an orbital radius of 3 mm until an OD_600_ of 0.5-0.8 was reached. OD_600_ was measured using a Thermo Scientific^™^ BioMate^™^ 160 UV-Vis Spectrophotometer. At this point, the pathway plasmids were fully induced with addition of corresponding inducer (1 mM IPTG, 1 mM vanillate, or 0.2 nM aTc), and the metabolite of interest was supplemented at this time. Cultures were incubated over 24 h at 37 °C with sampling and metabolite concentration measured via supernatant sampling and submission to HPLC.

For pN-Phe synthesis testing, cultures were inoculated with overnight culture grown in LBL-glucose media. Cultures were inoculated at 100x dilution from confluent overnight culture in 50 mL of the corresponding media with appropriate antibiotics in 250 mL baffled shake flasks and grown at 37 °C at 250 RPM. Expression vectors were fully induced at OD_600_ 0.5-0.8, and then were cultured at 30 °C. Synthesis of metabolites was quantified via supernatant sampling over 24 h and analysis by HPLC. Compound confirmation was performed by UPLC-MS.

### 2.4. Overexpression and purification of *N-*oxygenases

A strain of *E. coli* BL21 (DE3) harboring a pZE plasmid encoding expression of an *N-*oxygenase with a hexahistidine tag at either the N-terminus or C-terminus (NB01, NB02, NB03, or NB38) was inoculated from frozen stocks and grown to confluence overnight in 5 mL LBL containing kanamycin. Confluent cultures were used to inoculate 400 mL of experimental culture of LBL supplemented with kanamycin. The culture was incubated at 37 °C until an OD_600_ of 0.5-0.8 was reached while in a shaking incubator at 250 RPM. *N-*oxygenase expression was induced by addition of anhydrotetracycline (0.2 nM) and cultures were incubated at 30 °C for 5 h. Cultures were then grown at 20 °C for an additional 18 h. Cells were centrifuged using an Avanti J-15R refrigerated Beckman Coulter centrifuge at 4 °C at 4,000 *g* for 15 min. Supernatant was then aspirated and pellets were resuspended in 8 mL of lysis buffer (25 mM HEPES, 10 mM imidazole, 300 mM NaCl, 10% glycerol, pH 7.4) and disrupted via sonication using a QSonica Q125 sonicator with cycles of 5 s at 75% amplitude and 10 s off for 5 minutes. The lysate was distributed into microcentrifuge tubes and centrifuged for 1 h at 18,213X*g* at 4 °C. The protein-containing supernatant was then removed and loaded into a HisTrap Ni-NTA column using an ÄKTA Pure GE FPLC system. Protein was washed with 3 column volumes (CV) at 60 mM imidazole and 4 CV at 90 mM imidazole. *N-*oxygenase was eluted in 250 mM imidazole in 1.5 mL fractions. Selected fractions were denatured in Lamelli SDS reducing sample buffer (62.5 mM Tris-HCl, 1.5% SDS, 8.3% glycerol, 1.5% beta-mercaptoethanol, 0.005% bromophenol blue) for 10 minutes at 95 °C and subsequently run on an SDS-PAGE gel with a Thermo Scientific PageRuler™ Prestained Plus ladder to identify protein containing fractions and confirm their size. The *N-*oxygenase containing fractions were combined applied to an Amicon column (10 kDa MWCO) and the buffer was diluted 1,000x into a 20 mM Tris pH 8.0, 5% glycerol buffer.

### 2.5. *N-*oxygenase expression testing

To test expression of the *N-*oxygenase ladder, 5 mL cultures of NB17-NB37 were inoculated in 5 mL cultures of LBL containing 50 μg/mL kanamycin and then grown at 37 °C until mid-exponential phase (OD = 0.5-0.8). At this time, cultures were induced via addition of 0.2 nM aTc and then grown at 30 °C for 5 h before growing at 20 °C for an additional 18 h. After this time, 1 mL of cells was mixed with 0.05 mL of glass beads and then vortexed using a Vortex Genie 2 for 15 minutes. After this time, the lysate was centrifuged at 18,213 *g* at 4 °C for 30 minutes. Lysate was denatured as described in Section 2.5 and then subsequently run on an SDS-PAGE gel with an Invitrogen BenchMark^™^ His-tagged Protein Standard ladder and then analyzed via western blot with an HRP-conjugated 6*His His-Tag Mouse McAB primary antibody. The blot was visualized using an Amersham ECL Prime chemiluminescent detection reagent.

### 2.6. *In vitro N-*oxygenase activity assay

Reactions were performed in 1 mL volumes consisting of 25 mM HEPES pH 7.0, 25 mM NaCl, and 1.5% H_2_O_2_ with 1 mM pA-Phe or pA-Pyr. The reaction mixture was incubated for 3 h at 25 °C with 10 µM purified *N-*oxygenase, following which protein was removed by filtering through a 10 K Amicon centrifugal filter unit. The eluent was then analyzed by HPLC. pN-Phe production was further confirmed via UPLC-MS.

### 2.7. HPLC Analysis

Metabolites of interest were quantified via high-performance liquid chromatography (HPLC) using an Agilent 1260 Infinity model equipped with a Zorbax Eclipse Plus-C18 column. To quantify amine containing metabolites, an initial mobile phase of solvent A/B = 100/0 was used (solvent A, 20 mM potassium phosphate, pH 7.0; solvent B, acetonitrile/water at 50/50) and maintained for 7 min. A gradient elution was performed (A/B) with: gradient from 100/0 to 50/50 for 7-17 min, gradient from 50/50 to 100/00 for 17-18 min, equilibration at 100/0 for 18-22 min. A flow rate of 0.5 mL min^-1^ was maintained and absorption was monitored at 210 and 320 nm. To quantify nitro-group containing metabolites, we used an initial mobile phase of solvent A/B = 100/0 (solvent A, water, 0.1% trifluoroacetic acid; solvent B, acetonitrile, 0.1% trifluoroacetic acid) and maintained for 5 min. A gradient elution (A/B) was then performed with: gradient from 100/0 to 95/5 over 5-7 min, gradient from 95/5 to 90/10 over 7-10 min, gradient from 90/10 to 80/20 over 10-16 min, gradient from 80/20 to 70/30 over 16-19 min, gradient from 70/30 to 0/100 over 19-21 min, 0/100 over 21-23 min, gradient from 0/100 to 100/0 over 23-24 min, and equilibration at 100/0 over 24-25 min. The nitro product quantifying method used flow rate of 0.5 mL min^-1^ and monitored absorption at 280 and 320 nm.

### 2.8. Mass Spectrometry

Mass spectrometry (MS) measurements for small molecule metabolites were submitted to a Waters Acquity UPLC H-Class coupled to a single quadrupole mass detector 2 (SQD2) with an electrospray ionization source. Metabolite compounds were analyzed using a Waters Cortecs UPLC C18 column with an initial mobile phase of solvent A/B = 95/5 (solvent A, water, 0.1% formic acid; solvent B, acetonitrile, 0.1% formic acid) with a gradient elution from (A/B) 95/5 to 5/95 over 5 min. Flow rate was maintained at 0.5 mL min^-1^. For samples collected from *E. coli* growth cultures, an initial submission to an Agilent 1100 series HPLC system with a Zorbax Eclipse Plus C18 column was used to collect pN-Phe elution peaks for enhanced MS resolution. A 100 uL injection was made with an initial mobile phase of solvent A/B = 95/5 (solvent A, water, 0.1% trifluoroacetic acid; solvent B, acetonitrile, 0.1% trifluoroacetic acid) and maintained for 1 min. A gradient elution was then performed (A/B) with: gradient from 95/5 to 50/50 over 1-24 min, gradient from 50/50 to 95/5 over 24-25 min, equilibration at 95/5 for 25-27 min. Flow rate was 1 mL/min and metabolites were tracked at 270 nm. pN-Phe elution was identified at 7.20 min using a chemical standard and this peak was collected for submission to UPLC-MS.

## 3. Results

### 3.1. Pathway Metabolite Stability and Toxicity

To initially determine whether the nitroaromatic compounds and their amine precursors in our proposed pathway were non-toxic, we performed toxicity testing in a 96 deep-well plate format. Here, we supplemented 1 mM of our envisioned pathway metabolites (pA-Phe, pA-Pyr, pN-Phe, and pN-Pyr) into cultures of *E. coli* MG1655 (DE3) at inoculation in LBL media with addition of 5% DMSO to aid metabolite solubility. We grew cultures and monitored cell density, observing minimal effect of metabolite addition on doubling time with the exception of the metabolite pN-Pyr, which increased doubling time from 27.4 ± 0.1 min to 54.8 ± 0.6 min (**Fig. 2A**). Fortunately, neither the aromatic amines nor pN-Phe exhibited notable toxicity.

We then performed stability testing in a similar manner, with candidate metabolites added at 0.5 mM concentration at inoculation for cultivation at 37 °C (**Fig. 2B**). After 24 h, pN-Phe concentration remained at 71 ± 12% of its initial value, indicating promising stability, whereas only 32 ± 9% of the initial pN-Pyr supplied remained. We hypothesized that pN-Pyr may be a substrate of native *E. coli* aminotransferases given previously published *in vitro* data on activity of the aminotransferase TyrB (Hayashi et al., 1993; Onuffer et al., 1995). This possible cause of instability would be convenient for our pathway design as the final pathway step would be catalyzed natively. We performed an additional experiment to monitor pN-Pyr and pN-Phe concentrations simultaneously after supplementation of 0.25 mM pN-Pyr to *E. coli* MG1655 (DE3) at inoculation (**Fig. 2C**). Here, we found that 75 ± 13% of supplemented pN-Pyr was converted to pN-Phe after 24 h, confirming that endogenous *E. coli* enzymes will convert pN-Pyr to pN-Phe.

### 3.2. Initial *N-*oxygenase Characterization

Given that *E. coli* synthesis of pA-Phe and pA-Pyr has previously been established, we sought to identify an *N-*oxygenase that can fully oxidize pA-Phe or pA-Pyr. There are four distinct types of *N*-oxygenase that have been characterized to date (Nóbile et al., 2021): (i) Flavin-dependent monooxygenases, (ii) Heme-oxygenase-like diiron oxygenases, (iii) Rieske-type *N-*oxygenases, and (iv) Non-heme diiron monooxygenases. Investigation into flavin-dependent monooxygenases has shown that they commonly act on amine-containing sugars (Huynh et al., 2020; Thoden et al., 2013). Additionally, studies on the substrate scope of heme-oxygenase-like diiron oxygenases have been fairly limited (imidazole and N^ω^-methyl-l-arginine) (Hedges and Ryan, 2019; Ng et al., 2019). While the Rieske-type *N-*oxygenase PrnD has previously demonstrated activity on pA-Phe, reconstitution of Rieske *N-*oxygenases *in vivo* has been challenging in *E. coli* due to limits in iron-sulfur cluster incorporation into enzymatic cores (Tiwari et al., 2011). Previous studies have demonstrated the activity of the characterized non-heme diiron monooxygenase-type *N-*oxygenases AurF (Winkler and Hertweck, 2005) and CmlI (Lu et al., 2012) *in vivo*, but neither have been shown to effectively convert pA-Phe to pN-Phe (Chanco et al., 2014). More recent work identified putative diiron monooxygenase ObaC in the synthesis of obafluorin. In that work, the activity of ObaC on pA-Pyr was indirectly demonstrated via a colorimetric assay wherein NADH oxidation led to color shift in the presence of phenazine methosulfate, iron(II) sulfate and ObaC (Schaffer et al., 2017). Additional combination of ObaC with ObaH and ObaG identified a nitro-containing final product via mass spectrometry, but confirmation of full oxidation of pA-Pyr to pN-Pyr was not measured via mass spectrometry. To confirm ObaC activity on pA-Pyr and test activity on pA-Phe, we prepared three ObaC fusions (ObaC-C_term_His_6x_, N_term_His_6x_-ObaC, and N_term_βgal-ObaC-C_term_His_6x_ [βgal-ObaC]) and performed Ni-NTA chromatography. We isolated proteins of correct size for NtermHis6x-ObaC and βgal-ObaC, but were unable to isolate ObaC-CtermHis6x. We then performed in vitro characterization on N_term_His_6x_-ObaC and βgal-ObaC using H_2_O_2_ as a reductant to recycle the enzyme’s diiron core (Chanco et al., 2014). Using 10 µM purified enzymes, we ran reactions containing 1.5% H_2_O_2_ over 3 h with 1 mM of either pA-Pyr or pA-Phe. Our initial results indicated that N_term_His_6x_-ObaC was unable to achieve sufficient activity on pA-Pyr or pA-Phe for measurement on HPLC. However, reactions run with pA-Pyr and pA-Phe using βgal-ObaC obtained 127 ± 6 µM pN-Pyr and 105 ± 15 µM pN-Phe, respectively (**Fig. 3A**). Using UPLC-MS analysis, the pA-Phe supplemented experiment was confirmed to produce pN-Phe.

To validate ObaC activity *in vivo*, we performed a supplementation experiment with an MG1655 strain harboring a pZE plasmid vector bearing the *obaC* gene. Here, we grew cultures to mid-exponential phase, and then supplemented cultures with either pA-Pyr or pA-Phe and induced ObaC expression. These experiments resulted in extracellular production of 31 ± 1 µM pN-Phe with pA-Pyr supplementation or 240 ± 20 µM pN-Phe with pA-Phe supplementation after 24 h (**Fig. 3B**), demonstrating that ObaC could act on either pathway intermediate. Our low titers suggest that pA-Pyr or pA-Phe may be transport-limited upon supplementation and that pA-Phe may enter cells more easily than pA-Pyr.

### 3.3. Implementation of a *de novo* pN-Phe synthesis pathway

Given demonstrated activity of ObaC on desired aromatic amines and native conversion of pN-Pyr to pN-Phe, we then sought to confirm synthesis of pA-Phe using the PapABC enzymes. We chose to utilize the *papABC* genes from *S. venezuelae* rather than homologues in *P. fluorescens* given results from a previous study that suggested that the *S. venezuelae* enzymes result in greater pA-Phe titer (Chen et al., 2018). To start, we obtained the *papABC* operon from the Mehl Lab in the pLASC-lppPW plasmid and then moved these into an IPTG inducible pCola plasmid under the control of an IPTG-inducible T7-promoter system. Initial shake flask testing of this construct in *E. coli* MG1655(DE3) using M9-glucose minimal media resulted in low 24h post-induction pA-Phe titers of <10 µM (**Fig. 4A**, NB06). Expression of the feedback-resistant DAHP aldolase AroG*, which is known to enhance shikimate pathway flux (Kunjapur et al., 2014), was tested using an IPTG-inducible pACYC plasmid under the control T7-promoter system with pCola-papABC (NB07). This strain resulted in significant 24 h post-induction pA-Phe titer of 624 ± 79 µM.

**Figure 4.**
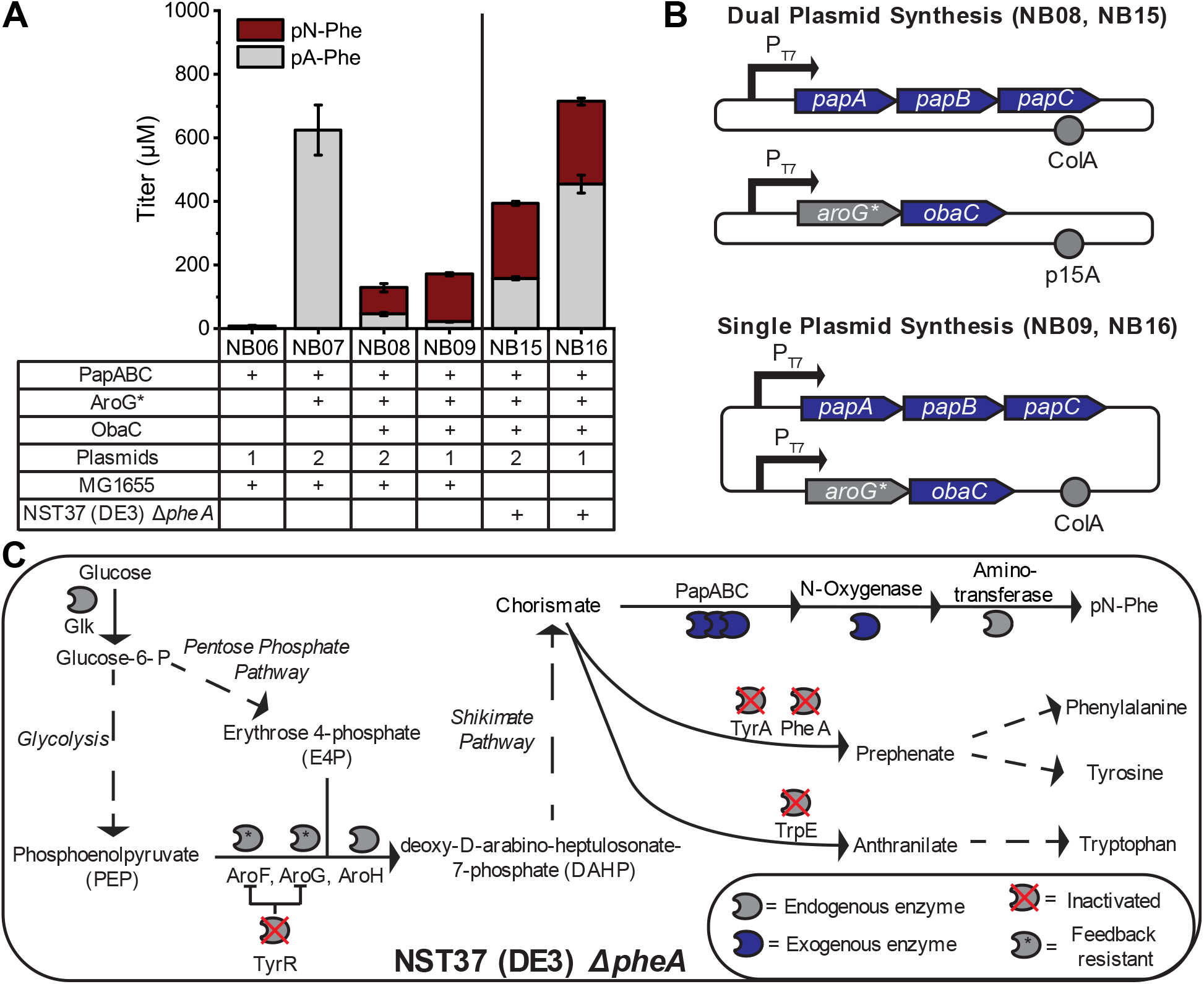
*De novo* synthesis of pN-Phe in *E. coli*. (A) Titer of pA-Phe and pN-Phe in different strains expressing pathway enzymes following 24 h of fermentation in M9-glucose (1.5%) minimal media Sample size is *n*=3 using biological replicates. Data shown are mean ± standard deviation. (B) Depiction of the different arrangement of pathway plasmids for the synthesis of pN-Phe. (C) Depiction of the E. coli NST37(DE3) ΔPheA strain engineered for enhanced pN-Phe titer through chorismate. Inactivated enzymes are shown with a red cross and feedback-resistant versions of enzymes are shown with a star (*). GIK, glucokinase; AroG, phenylalanine-sensitive Phospho-2-dehydro-3-deoxyheptonate (DAHP) synthase; AroF, tyrosine-sensitive DAHP synthase; AroH, tryptophan-sensitive DAHP synthase; TyrA and PheA, TyrA and PheA subunits of the chorismate mutase; TrpE, anthranilate synthase component I; TyrR, transcriptional regulatory protein.

We then incorporated the *N-*oxygenase ObaC into the pathway in a format using two plasmids: (i) a pACYC vector expressing AroG* and ObaC in an operon and (ii) the pCola vector expressing the PapABC operon (NB08). Here we first obtained *de novo* biosynthesis of pN-Phe, with a titer of 83 ± 13 µM pN-Phe and 46 ± 5 µM pA-Phe (**Fig. 4A**). We confirmed that the product mass of the peak of interest corresponded to pN-Phe by UPLC-MS analysis of a second NB08 fermentation in minimal media supplemented with glucose. Then, we cloned a single pCola vector expressing AroG* and ObaC in an IPTG-inducible operon and PapABC on an additional IPTG-inducible operon and expressed this in MG1655 (DE3) (NB09) (**Fig. 4B**). This strain demonstrated superior pN-Phe synthesis achieving a higher 24h post-induction titer of pN-Phe of 149 ± 5 µM in addition to a higher titer of pA-Phe of 153 ± 22 µM.

To further enhance titer, we purchased a commercially available phenylalanine overproducer strain and performed several modifications to improve its suitability for pN-Phe biosynthesis. We inactivated the expression of the initial enzyme which converts chorismate flux to phenylalanine, PheA (Rodriguez et al., 2014). We additionally performed genomic integration of the T7-polymerase and *lacI* and *lacZ* genes for use with our plasmid setup, creating the NST37 (DE3) *ΔpheA* strain. This strain is unable to express functional proteins for the synthesis of all three standard aromatic amino acids (TyrA, TrpE, PheA) and a shikimate pathway regulator (TyrR). Furthermore, the strain expresses feedback inhibition enzyme variants (AroF* and AroG*) to enhance shikimate pathway flux, which should result in greater accumulation of chorismate (**Fig. 4C**). Using this modified strain, we transformed both the two-plasmid system as previously described (NB15) and the single plasmid system (NB16) and performed shake-flask fermentation over 24 h in M9-glucose minimal media. Here, we achieved our highest measured titer of 260 ± 11 µM using the single plasmid system. Despite promising pN-Phe titer, in the case of the single plasmid system, the unconverted pA-Phe titer remained quite high at 454 ± 28 µM.

### 3.4. Broad *N-*oxygenase screening and pathway implementation

Previous studies using genetic code expansion for site-specific translation of pN-Phe into proteins have supplemented 1 mM of the nsAA into culture to achieve incorporation (Grünewald et al., 2008; Tsao et al., 2006), but lower concentrations of nsAA were not tested. While the obtained titer of approximately 250 µM may be sufficient for incorporation, high accumulation of pA-Phe could result in off-target nsAA incorporation if a strain were to perform biosynthesis and site-specific nsAA incorporation. Additionally, poor selectivity of pN-Phe biosynthesis would complicate downstream isolation of the small molecule product if that option were to be pursued. Thus, to improve product titer and make pN-Phe the dominant nsAA product, we sought to identify an *N-*oxygenase enzyme with improved activity on pA-Phe. Previously characterized non-heme diiron monooxygenases have demonstrated activity on diverse aromatic amines and compatibility with expression in *E. coli*, but ObaC is the only diiron monooxygenase thus far to exhibit full oxidation of pA-Phe to pN-Phe. To assess the broader space of *N-*oxygenase enzymes, we used NCBI BLAST to obtain the 1000 most closely related sequences as measured by BLASTP alignment score from four characterized diiron monooxygenase-type *N-*oxygenases with activity on aromatic amines: AurF, CmlI, AzoC (Guo et al., 2019), and ObaC. After deleting duplicate sequences, we obtained 2134 unique sequences which we then submitted to the Enzyme Function Initiative-Enzyme Similarity Tool (EFI-EST) (Gerlt et al., 2015) to generate a sequence similarity network (SSN). Sequences exhibiting greater than 95% similarity were grouped into single nodes, resulting in 775 unique nodes and a minimum alignment score of 100 was selected for node edges. We visualized the SSN using Cytoscape (**Fig. 5**).

**Figure 5.**
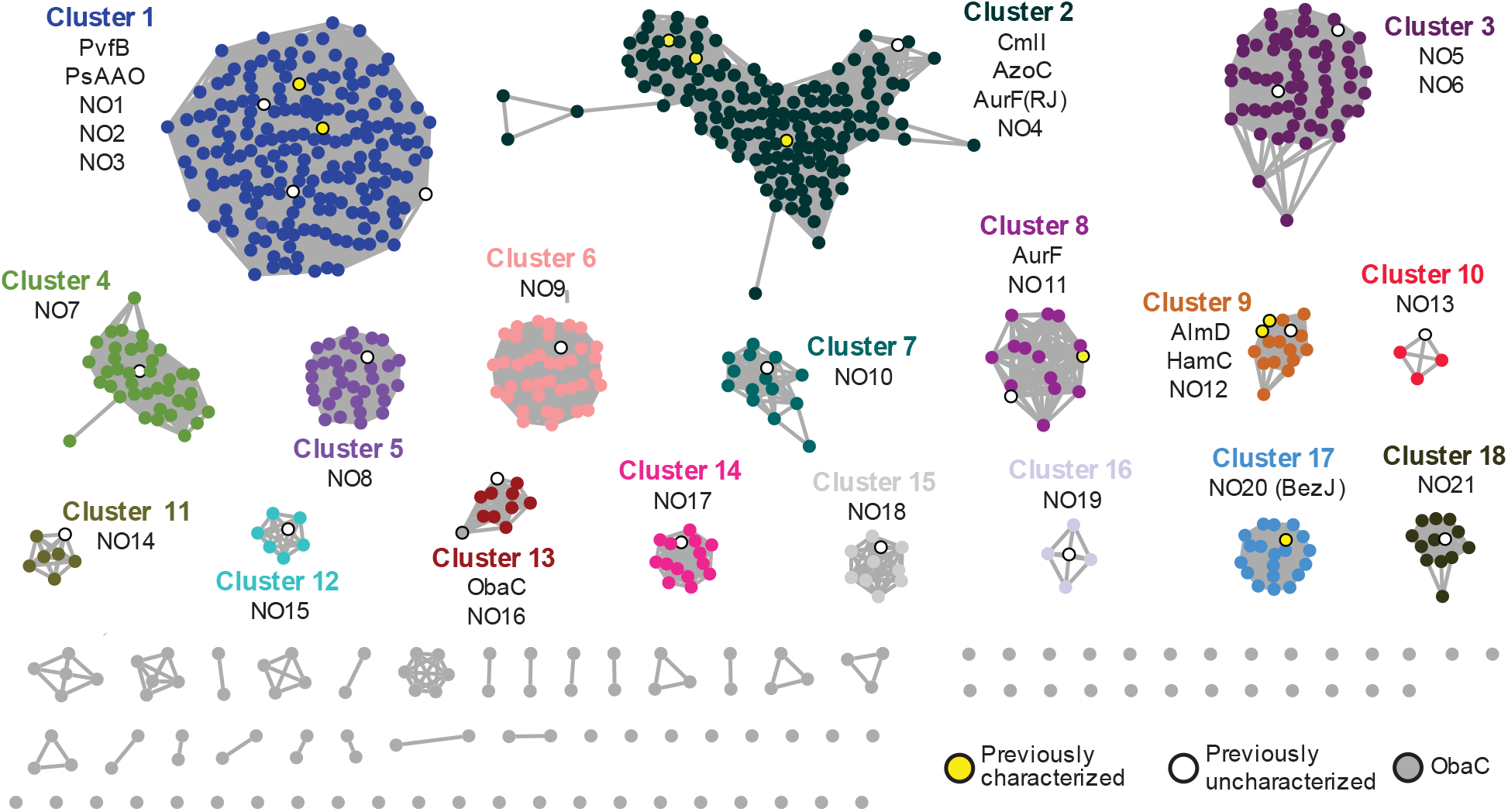
Sequence similarity network generated using 2134 unique putative diiron monooxygenase sequences determined from NCBI BLAST of the *N-*oxygenases AurF, CmlI, AzoC, and ObaC represented as 775 unique nodes. Edges are drawn between nodes with minimum alignment score of 100. Previously characterized *N-*oxygenases are highlighted in yellow (PvfB (Kretsch et al., 2018), PsAAO (Platter et al., 2011), CmlI (Knoot et al., 2016; Lu et al., 2012), AzoC (Guo et al., 2019), AurF(RJ) (Indest et al., 2015), AurF (Seong Choi et al., 2008), AlmD (Cortina et al., 2011), HamC (Jenul et al., 2018), and BezJ (Tsutsumi et al., 2018)), putative *N-*oxygenases cloned and tested in this study are highlighted in white, and ObaC is shown in gray with black outline.

To test a diverse set of *N-*oxygenase enzymes, we selected twenty-one distinct sequences from 18 different clusters to obtain diversity in the diiron monooxygenase sequence space from those currently characterized and ordered gene fragments to clone these into a pZE expression vector with a C-terminal hexahistidine tag. All but one of these *N-*oxygenases (NO20, also known as BezJ) (Tsutsumi et al., 2018)) were previously uncharacterized. We initially tested the expression of these proteins in *E. coli* by transforming them into MG1655(DE3) for subsequent overnight expression. Analysis of the lysate using anti-His western blotting confirmed the soluble synthesis of all the *N-*oxygenases except NO6, NO13, NO18, and NO20. We then tested the *N-*oxygenases in MG1655(DE3) by growing them in LBL media to mid-exponential phase, at which time, we supplemented cultures with 1 mM pA-Phe and induced *N-*oxygenase expression. Fermentation with metabolically active cells for 24 h at 30 °C revealed that only one of the additional *N-*oxygenases resulted in pN-Phe production apart from ObaC. This *N*-oxygenase was NO16 from *Pseudomonas sp. EMN2*, which exhibited greater than twice the pN-Phe production in the initial screen (**Table 1**). Previous reports of *para*-aminobenzoic acid testing have noted activity in enzymes across different clusters (PsAAO (Platter et al., 2011), AurF (Seong Choi et al., 2008), and HamC (Jenul et al., 2018)), which we did not observe for pA-Phe. Following the initial *in vivo* screen, we purified NO16 with a C-terminal hexahistidine tag and performed a 3 h *in vitro* reaction containing 1.5% H_2_O_2_ with 1 mM of either pA-Pyr or pA-Phe. In comparison to ObaC, we obtained similar titer of pN-Pyr in this case (120 ± 3 µM final concentration pN-Pyr) and a two-fold increase in the titer of pN-Phe (196 ± 23 µM final concentration of pN-Phe) when NO16 is used as catalyst (**Fig. 6A**).

**Table 1.** *In vivo* screening of *N*-oxygenase activity on pA-Phe via supplementation testing of 1 mM pA-Phe in cultures expressing *N-*oxygenases with C-terminal hexahistidine tags via measurement of pN-Phe titer after 24 h fermentation at 37 °C. ObaC was tested without a hexahistidine tag.

| *N*-oxygenase | pN-Phe Titer<br>(μM) |
| --- | --- |
| ObaC | 45 ± 19 |
| NO6, NO13, NO18,<br>NO20 | No protein<br>detected |
| NO1-5, NO7-12, NO14,<br>NO15, NO17, NO19,<br>NO21 | No product<br>detected |
| NO16 | 105 ± 39 |

**Figure 6.**
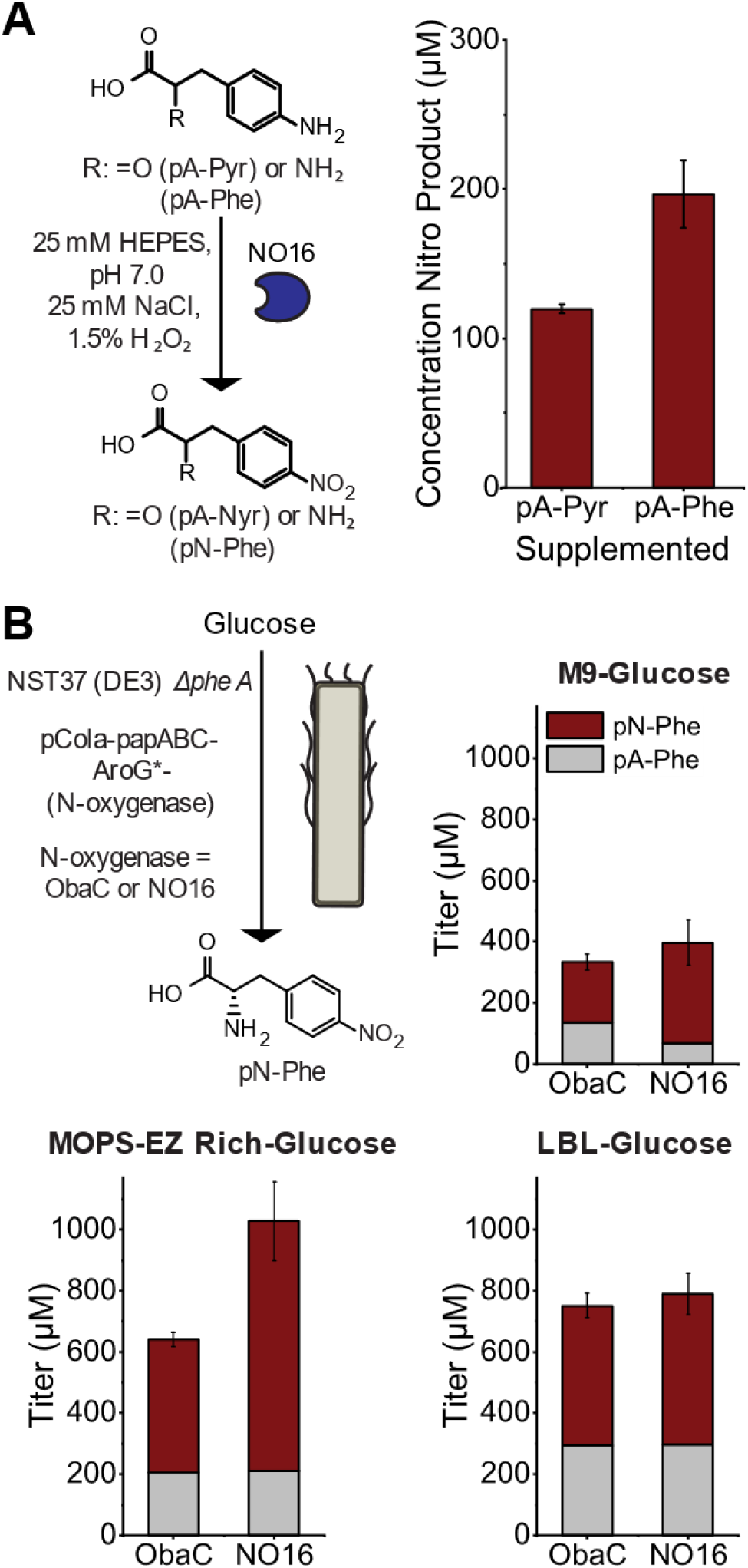
Evaluation of the *N-*oxygenase NO16 in the synthesis of pN-Phe. (A) In vitro investigation of NO16 activity on pA-Phe and pA-Pyr. (B) Media comparison of the pN-Phe and pA-Phe titers using the NST(DE3) *ΔpheA* strain of *E. coli* transformed with either pCola-papABC-AroG*-ObaC (strain NB16) or pCola-papABC-AroG*-NO16 (strain NB39) in either M9-glucose, MOPS-EZ rich-glucose, or LBL-glucose media.

Then, we sought to compare the performance of NO16 and ObaC within our *de novo* biosynthesis pathway for pN-Phe. To start, we cloned an additional pN-Phe synthesis vector with NO16 in place of ObaC in the operon with *aroG\**. We then evaluated the 24 h titer in M9-glucose minimal media of the pCola-papABC-AroG*-NO16 construct in the NST37 (DE3) Δ*pheA* strain in comparison to the NST37 (DE3) Δ*pheA* strain expressing the pCola-papABC-AroG*-ObaC plasmid. Here, we saw enhanced pN-Phe titer in the strain expressing NO16 (330 ± 75 µM) as compared to the strain expressing ObaC (197 ± 25 µM) (**Fig. 6B**).

With this new *N*-oxygenase in hand, we finally evaluated the pN-Phe titer observed at 24 h that we could obtain in richer growth media at shake flask scale. Using MOPS EZ rich glucose defined media, we saw the highest pN-Phe titers of 820 ± 130 µM when using NO16 in the pCola vector compared to 435 ± 23 µM when using ObaC (**Fig. 6B**). In richer LBL-glucose media, we saw lower titers of pN-Phe titers when using NO16 (492 ± 68 µM) and comparable titers to MOPS EZ rich when using ObaC (456 ± 40 µM). The better performance of the rich-defined media using NO16 was additionally achieved with a lower final cell density of approximately half that of LBL media.

## 4. Conclusion

This work is the first to demonstrate *de novo* biosynthesis of a non-naturally occurring nsAA. In addition, prior to this work *N-*oxygenases had not been harnessed in biosynthetic pathways for novel nitro-product biosynthesis. The potential to safely synthesize nitroaromatic compounds from renewable feedstocks is valuable, and further studies into nitroaromatic biosynthesis using *N-*oxygenases are warranted. Here, we demonstrated the biosynthesis of the nitroaromatic amino acid pN-Phe in titers of 330 ± 75 µM pN-Phe from M9-minimal media and 820 ± 130 µM from a MOPS-EZ rich defined media, with pN-Phe representing the dominant heterologous metabolite. Utilization of this pathway with methods of genetic code expansion could enable the use of pN-Phe within proteins for *in situ* applications and further broaden its potential use case.

## 6. Author Statements

A.M.K. conceived and supervised the study; N.D.B. designed and performed all experiments, analyzed data, prepared figures, and wrote the manuscript; M.L. aided with molecular cloning; SS cloned the N-oxygenase library and confirmed expression.

## 8. Acknowledgements

We are grateful to Dr. PapaNii Asare-Okai of the University of Delaware Chemistry Mass Spectrometry Facility for assistance with metabolite LC-MS. We acknowledge support from the following funding sources: The National Science Foundation (NSF CBET #2032243), University of Delaware Start-Up Funds, the Mort Collins Foundation, and minor research support as part of the Center for Plastics Innovation, an Energy Frontier Research Center funded by the U.S. Department of Energy (DOE), Office of Science, Basic Energy Sciences, under Award No. # DE-SC0021166 (mass spectrometry analysis, *N*-oxygenase screening, manuscript preparation).

